# BmCncC/keap1-pathway is involved in high-temperature induced metamorphosis regulation of silkworm, *Bombyx mori*

**DOI:** 10.1101/458620

**Authors:** Jinxin Li, Tingting Mao, Zhengting Lu, Mengxue Li, Zhengting Lu, Jianwei Qu, Yilong Fang, Jian Chen, Hui Wang, Xiaoyu Cheng, Hu Jiahuan, Yu Liu, Zhang Ziyan, Gu Zhiya, Li Fanchi, Bing Li

**Affiliations:** School of Basic Medicine and Biological Sciences, Soochow University, Suzhou, Jiangsu 215123, P.R. China; Sericulture Institute of Soochow University, Suzhou, Jiangsu 215123, P.R. China

## Abstract

The global warming has affected the growth, development and reproduction of insects. However, the molecular mechanism of high temperature stress-mediated metamorphosis regulation of lepidopteran insect has not been elucidated. In this study, the relationship between the insect developmental process and endogenous hormone level was investigated under high temperature (36 ° C) stress in *Bombyx mori* (*B. mori*). The results showed that the duration of 5^th^ instar larvae were shortened by 28 ± 2 h, and the content of 20E was up-regulated significantly after 72 h of high temperature treatment, while the transcription levels of 20E response genes *E93, Br-C, USP, E75* were up-regulated 1.35, 1.25, 1.28, and 1.27-fold, respectively. The high temperature treatment promoted the phosphorylation level of Akt and the downstream BmCncC/keap1 pathway was activated, the transcription levels of 20E synthesis-related genes *cyp302a1, cyp306a1, cyp314a1* and *cyp315a1* were up-regulated by 1.12, 1.51, 2.17 and 1.23-fold, respectively. After treatment with double stranded RNA of BmCncC (dsBmCncC) in BmN cells, the transcription levels of *cyp302a1* and *cyp306a1* were significantly decreased, whereas up-regulated by 2.15 and 1.31-fold, respectively, after treatment with CncC activator Curcumin. These results suggested that BmCncC/keap1-mediated P450 genes (*cyp302a1, cyp306a1*) expression resulted in the changes of endogenous hormone level, which played an important role in the regulation of metamorphosis under high temperature stress. Studies provide novel clues for understanding the CncC/keap1 pathway-mediated metamorphosis regulation mechanism in insects.

**Author Summary:** Mammalian nuclear transcription factor Nrf2 (NF-E2-related factor 2) plays an important role in the stress response of cells. CncC is a homolog of mammalian Nrf2 in insect, regulating the genes expression of insect antioxidant enzymes and cytochrome P450 detoxification enzyme. Evidence suggests that the CncC/Keap1 pathway also plays an important role in regulating insect development. Here, we investigated the regulatory mechanism between the CncC/Keap1 pathway and metabolism of silkworm hormones in Lepidoptera. We found that high temperature induction accelerated the development of silkworm, the ecdysone content and related metabolic genes in hemolymph were significantly up-regulated, the CncC/Keap1 pathway was activated, and the expression of *BmCncC* was significantly increased, indicating that the Cncc/Keap1 pathway plays an important role in this process. The expression of *cyp302a1* and *cyp306a1* was significantly decreased by RNA interference with *BmCncC*, which indicated that CncC in silkworm had a regulatory relationship with downstream 20E synthetic gene. In summary, the results indicate that the CncC/Keap1 pathway plays an important role in regulating hormone metabolism in silkworm, providing a basis for further study of the relationship between CncC/Keap1 pathway and development in insects.

## Introduction

The global warming resulted from greenhouse-effect adversely affects the ecosystems and survival of organisms, changing the living habitats and causing deterioration of the ecological environment [1, 2]. Insects are the most abundant species on earth and play an important role in the balance of ecosystems and the human agroforestry economy [3]. Insects are temperature-variable animals that are extremely sensitive to environmental changes. The high temperature environment directly affects the growth and development of insects, changing the biological characteristics, reproductive ability and life span [4,5,6]. Silkworm (*Bombyx mori*) belongs to the Lepidoptera and is an important economic insect [7, 8] which is susceptible to high temperature environment during rearing, resulting in reduction of survival rate, cocoon rate, and pupa yield [9,10]. The molecular mechanism of high temperature-mediated interaction between metamorphosis processes and endogenous hormone metabolism has not been elucidated.

The mammalian Nrf2 (nuclear factor erythroid 2 related factor 2)/Keap1 (Kelch-like ECH-associated protein 1) pathway regulates intracellular redox potential, metabolic detoxification, enhances cell resistance and delays aging [11,12]. Under stress conditions, Nrf2 induces the expression of antioxidant and detoxification enzyme genes by selectively binding to the antioxidant regulatory element (ARE) element in the promoter region to protect the cells against external stress [13, 14]. CncC (cap ‘n’ collar isoform-C) is the homolog of Nrf2 in insects, which plays an important role in metabolic detoxification and antioxidant enzyme genes expression [15]. The cytochrome P450 family genes in insects are mainly involved in detoxification of plant toxins and exogenous chemicals, which are related to insecticides resistance, and also play important role in biosynthesis of insect hormones such as 20E and pheromone et. [16]. Inhibition of CncC gene expression can change the developmental process of *Leptinotarsa decemlineata* [17], suggesting that CncC plays an important role in the insect metamorphosis. The P450 family genes in silkworm are divided into four subfamilies according to their homology: CYP2, CYP3, CYP4 and mitochondrial P450 [18], and the Hollween family genes *cyp302a1*, *cyp306a1*, *cyp314a1* and *cyp315a1* are mainly involved in 20E synthesis [19]. In silkworm, whether the P450 family Hollween genes are regulated by the BmCncC/keap1 pathway under high temperature stress have not been reported.

## Results

### Effects of HT stress on the development and vitality of silkworm

The results indicate that high temperature treatment resulted in smaller body size of larvae and transparent epidermis (Fig 1A), the characteristics and behavior of mature silkworm larvae were observed either. In addition, the larvae maturity time was advanced 28±2 h in average and mostly were concentrated around 144 h to 168 h (Fig 1C), indicating that high temperature treatment shortens the developmental duration of larvae. Furthermore, the vitality and pupation rate in high temperature group was reduced by 0.85 and 0.76-fold of the control group, respectively (Fig 1B and D), indicating that high temperature treatment reduced the vitality of the silkworm.

**Fig. 1.**
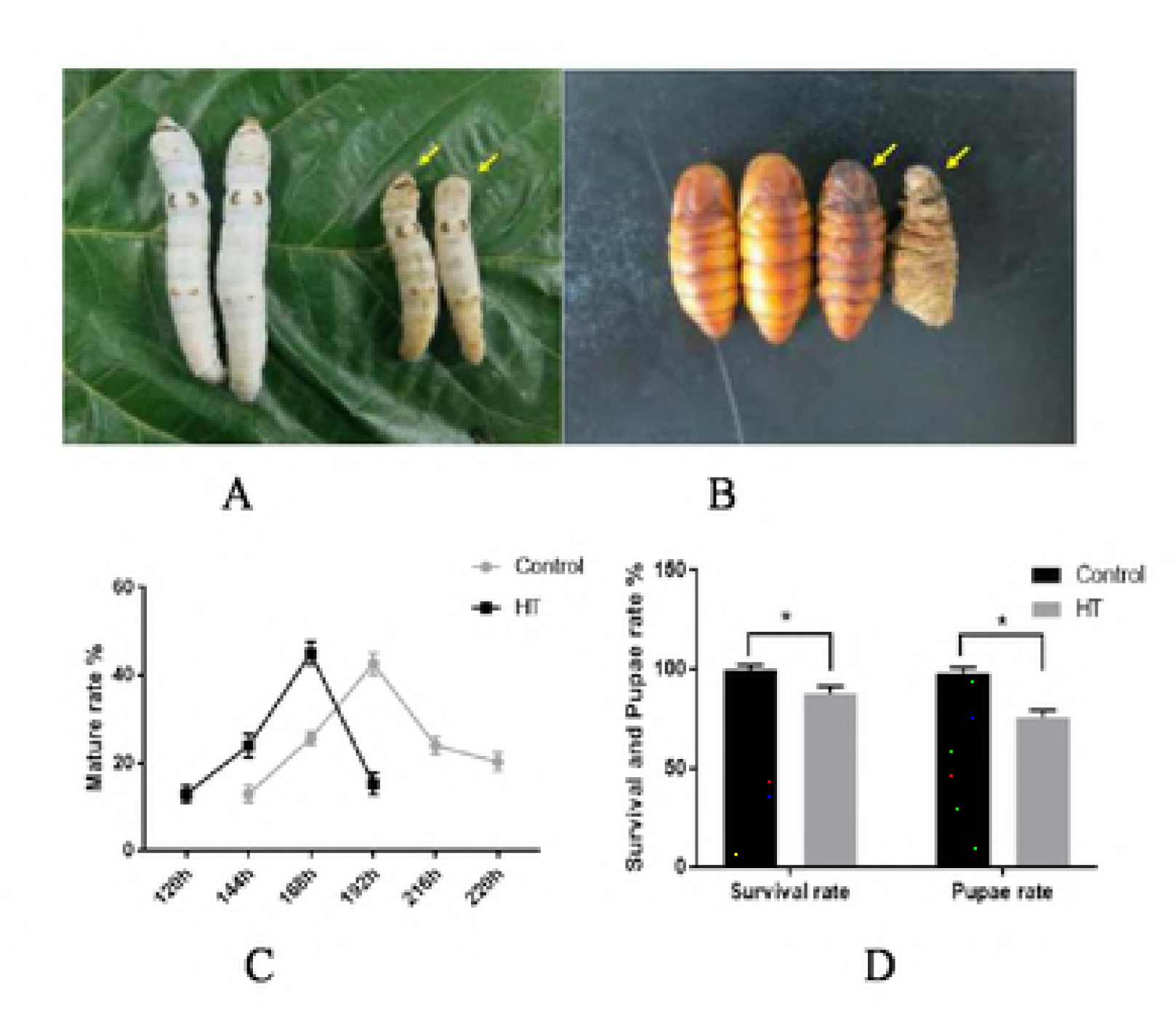
Effect of HT on the development and vitality of *B.mori*. A. Effect of HT on the larvae of silkworm. B. Effect of HT on the development of pupa. C. Effect of HT on the silkworm duration of developmental stage. D. Effect of HT on silkworm vitality and percentage of pupation. HT: High Tempature. The experiment was repeated three times independently. The results were shown to be the means values, and the siginificant differences were expressed by *(P≥0.05).

### The effect of HT stress on endogenous hormone and its metabolism-related genes expression

The content of JH in hemolymph was reduced by 0.63-fold at 72 h after high temperature treatment (Fig 2A), and the mRNA levels of the JH metabolism genes *Met*, *JH3* and *Kr-h1* were down-regulated by 0.75, 0.77, and 0.54-fold, respectively (Fig 2C). In addition, the content of 20E was increased by 1.86-fold after 72 h treatment, and the expression of 20E response genes *USP*, *Br-C*, *E75* and *E93* was up-regulated by 1.76, 1.35, 2.94 and 1.79-fold, respectively (Fig 2D). These results indicated that high temperature treatment changed the level of endogenous hormone in hemolymph.

### High temperature stress activated Akt/BmCncC/Keap1 pathway and induced expression of 20E-synthetic genes

The mRNA levels of BmCncC and Maf were up-regulated by 1.89 and 1.32-fold, and keap1 was down-regulated by 0.85-fold. The transcription levels of upstream genes PI3K and Akt was up-regulated significantly by 1.53 and 2.59-fold, respectively. Western blot analysis showed that the protein levels of Akt, p-Akt and BmCncC were up-regulated by 1.45, 1.56 and 1.32-fold, while the protein level of Keap1 was down-regulated by 0.76-fold, respectively. Furthermore, the mRNA levels of 20E-biosynthesis genes *cyp302a1*, *cyp306a1*, *cyp314a1*, and *cyp315a1* were significantly up-regulated, which was 1.12, 1.51, 2.17, and 1.23-fold, respectively (Fig 3A). The results indicated that high temperature treatment activated the Akt/BmCncC/keap1 pathway and induced the expression of downstream 20E biosynthesis genes.

### Regulation of downstream P450 genes by BmCncC

The mRNA levels of *cyp302a1* and *cyp306a1* in BmN cells were significantly down-regulated by 0.46 and 0.78-fold after treatment with dsBmCncC for 24 h. In contrast, after treatment with CncC activator Curcumin for 24 h, the mRNA levels of *cyp302a1* and *cyp306a1* were up-regulated by 2.15 and 1.31-fold, respectively (Fig 4). The transcriptional levels of *cyp314a1* and *cyp315a1* were non-significantly changed after treatment (Fig 4). These results indicated that BmCncC plays an important role in regulation of *cyp302a1* and *cyp306a1* genes.

## Discussion

The high temperature environment caused by climate warming resulted in the deterioration of the ecological environment and affects the growth and development of animals and plants [20]. The elevate in temperature can increase the annual generations of insects in the growing season, affecting the growth and development of insects [21]. Studies have shown that high temperature stress can reduce the vitality, body weight, and cocoon shell rate of *B. mori*, and promote the expression of stress-related genes and CSP (chemosensory protein) genes [10]. In this study, the larva epidermis became transparent and showed the characteristics of mature larva after high temperature induction (Fig 1A), the survival and pupation rate of silkworm were significantly decreased (Fig 1D), which was in consistent with previous studies [10]. BmCncC/keap1 pathway plays an important role in response to oxidative stress and regulation of antioxidative and detoxification genes expression [22]. Whether this pathway involved in metamorphosis processes and regulates ecdysone-synthesis P450 genes in silkworm have not been reported. We found that the duration of 5^th^ instar larva was shortened and the maturity time was advanced and relatively concentrated after high temperature induction (Fig 1C), indicating that the high temperature promoting the development of silkworm. We also investigated the relationship between the 20E synthesis-related P450 genes and BmCncC/keap1 pathway by alternating the endogenous hormone levels under high temperature induction.

The steroid hormone ecdysone and JH coordinated to regulate the development of insects in an opposite way. JH treatment can prolong the duration of larva and enlarge the body size [23]. In the study, the content of JH and the transcriptional levels of response genes *Met*, *JH3*, and *Kr-h1* were significantly down-regulated after high temperature treatment (Fig 2A and Fig 2C). The *JH3* and *Met* genes play an important role in the JH signaling cascade [24]. *Kr-h1* can maintain *B. mori* larval state by controlling 20E content mediated by inhibition of steroidogenic enzymes genes transcription [25]. The 20E content was increased significantly after high temperature treatment, and the transcriptional levels of 20E metabolism-related genes *E93*, *Br-C*, *USP* and *E7*5 were significantly up-regulated (Fig 2D). Previous studies shown that treatment silkworm with 20E at 5^th^ instar larva can accelerate the development and promote larvae maturation [26]. *E93* gene promotes the development of silkworm and induces the expression of downstream 20E response genes, including *Br-C*, *USP* and *E75*, that plays important role in 20E signal cascade [27]. The results suggest that high temperature stress alternated the normal level of endogenous hormone, and the significant up-regulation of 20E content and response-genes under high temperature induction maybe one of the reasons for the accelerated development of silkworm.

Hollween genes belongs to the insect P450 genes family play an important role in the synthesis of 20E [28]. In this study, the transcription levels of 20E-synthesis genes (*cyp302a1*, *cyp306a1*, *cyp314a1*, *cyp315a1*) were significantly up-regulated after high temperature induction (Fig 3A). Studies shown that treatment silkworm with exogenous substances (TiO_2_ NPs) promoted the expression of Hollween genes and increased the content of 20E [29], our results demonstrate that high temperature treatment promotes the anabolic level of 20E as well. In this study, high temperature treatment activated PI3K, resulting in phosphorylation of Akt and increased the expression of downstream BmCncC (Fig 3C), suggesting that the PI3K/Akt/BmCncC axis plays an important role in the up-regulation of Hollween genes expression under high temperature. Evidence suggests that CncC regulates the expression of the downstream P450 family genes [30]. Studies shown that CncC constitute a homodimer or a heterodimer with Maf to regulate the expression of the P450 genes [31]. The CncC and Maf complex regulates the expression of P450 genes by regulating the promoter activities of *CYP389B1* and *CYP392A28* genes in *Boisduval* (*Tetranychus cinnabarinus*) [32]. Inhibition of the CncC/keap1 pathway altered the metamorphosis process of the Colorado potato beetle (*Leptinotarsa decemlineata*), suggesting that the CncC/keap1 pathway plays an important role in developmental regulation [33]. Our study investigated whether BmCncC regulated the downstream 20E-synthesis P450 genes, found that the mRNA level of *cyp302a1* and *cyp306a1* was significantly down-regulated after inhibition of *BmCncC* (Fig 4), whereas significantly up-regulated after treatment with Curcumin, indicating that a regulatory relationship exists between *BmCncC* and downstream P450 (*cyp302a1 cyp306a1*) genes. Furthermore, the mRNA levels of *cyp314a1* and *cyp315a1* were not changed significantly after activation or inhibition of *BmCncC*, implying that there is an unknown pathway of regulation.

In summary, our study demonstrated that high temperature induction can accelerate the development of silkworm, elevate the content of 20E and promote related-genes expression, indicating a regulatory relationship between *BmCncC* and downstream 20E-biosynthetic genes, *cyp302a1* and *cyp306a1* in silkworm. Our results provided new clues for further studying of the high temperature impact on insect hormone metabolism and the regulatory relationship between *CncC* and P450 family genes.

## Methods

### Insects and treatment

The larvae of *B. mori* (Jingsong × Haoyue strain) maintained in our laboratory were reared on mulberry leaves under 12 h light/ 12 h dark conditions at 25 ± 1°C with 75-85% humidity. After three days of normal rearing, the 5^th^ instar silkworms were maintained at constant high temperature of 36 °C and 75 ± 5% humidity, and feed with fresh mulberry leaves.

### Cell culture and treatment

The BmN cells were maintained at 27°C in Grace insect medium, supplemented with 10% fetal bovine serum and 1% antibiotics. DsRNA of BmCncC was purchased from Shanghai GenePharma Co., Ltd, China. BmN cells were cultured in 12-well plates for 6 h and the medium was replaced with serum-free medium without antibiotics. Cells were transfected using 1μl (20 μM/μL) siRNA mixed with LipoHigh Liposome efficient transfection reagent (Sangon, Shanghai, China) and incubated for 8 h. Cells were cultured with dsRNA for another 24 h after replacement of medium. For Curcumin treatment (final concentration 10μM/ml), the cells were analyzed after 24 h treatment with Curcumin. All data are expressed as mean of 3 replicates.

### Elisa analysis

Ecdysone and juvenile hormone (JH) content were determined using a kit (MEIMIAN, Shanghai, China), refer to the instructions. The absorbance value was measured at wavelength of 450 nm to calculate the sample content. All data are expressed as mean of 3 replicates.

### Real time-quantitative PCR (qRT-PCR) analysis

Total RNA was extracted from silkworm fat body using RNA lysate (Takara, China). Primer sequences are shown in Table 1. Use *Actin3* as the internal reference gene. QRT-PCR was performed on a ViiA 7 System (ABI, Foster City, CA, USA). The reaction was in 20 μL volume. Amplification conditions were as follows: denaturation at 95°C for 1 min, followed by 45 cycles of 95°C for 5 s, 55°C for 10 s, and 72°C for 10 s. Data are expressed as the mean of three independent experiments ± SE (standard error).

**Table 1.**
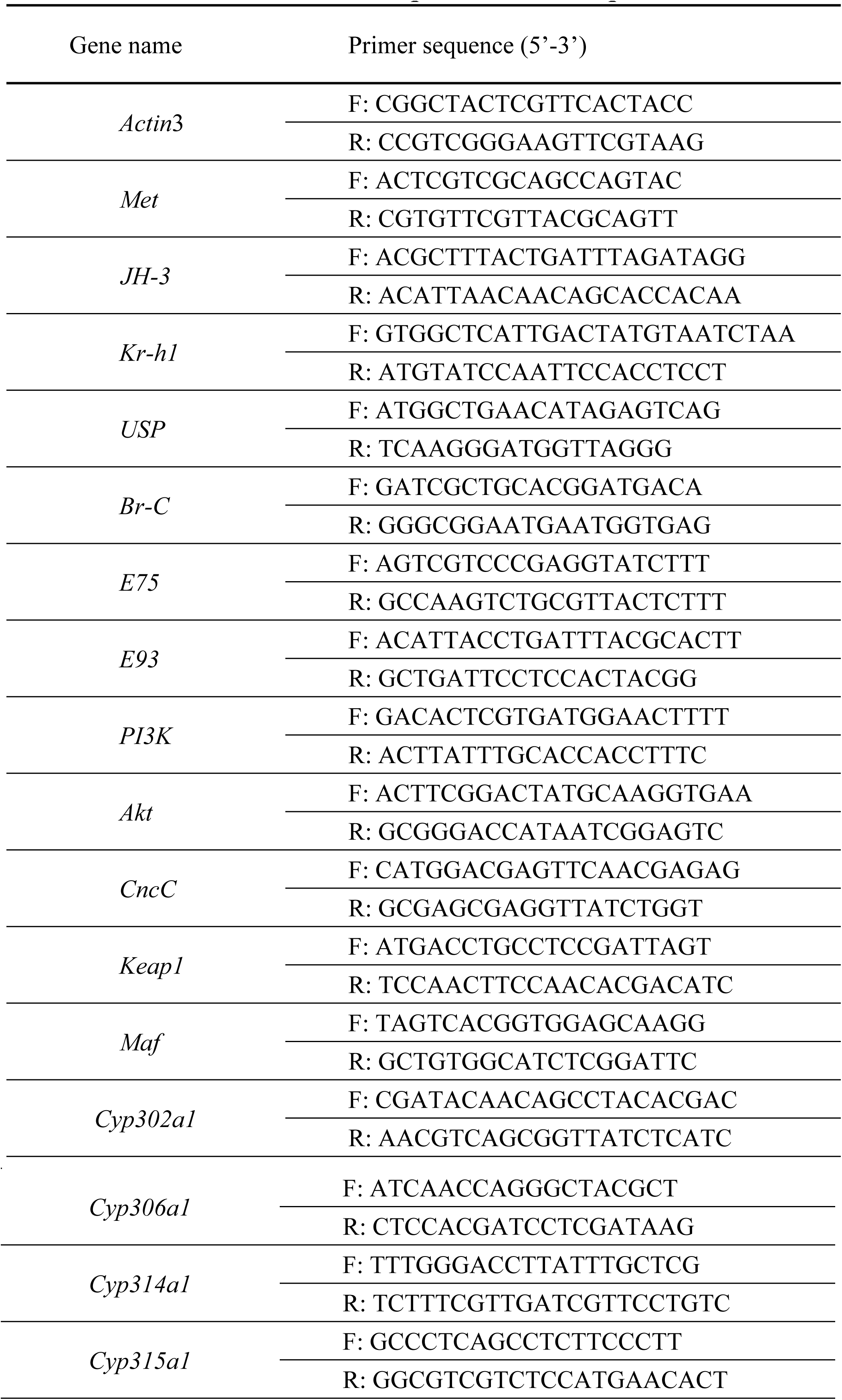
Primer sequences used in qRT-PCR

### Western blotting analysis

The samples of fat body of the control and treated groups were homogenized in lysis buffer supplemented with 1 mM PMSF. The samples were centrifuged at 12000 g for 10 min, and the supernatant was collected for analysis. The procedure was carried out according to the method before [26]. The Akt, p-Akt antibodies (CST, USA, 1:2000), polyclonal antibodies for BmCncC, Bmkeap1 (GenScript, Shanghai; 1:1500) were used as the primary antibody, and the HRP-conjugated goat anti-rabbit IgG (CST, USA, 1:2000) was used as the secondary antibody.

### Statistical analysis

All data are expressed as mean of 3 replicates. The differences in means between multiple sets of data were compared by one-way ANOVA. Dunnett’s test was performed when compared with the control. P <0.05 was considered significant difference.

## Acknowledgement

This study was supported by the China Agriculture Research System (CARS-18-ZJ0106), the State Key Laboratory of Integrated Management of Pest Insects and Rodents (Y852981303), the Natural Science Fund project in Jiangsu Province (BK20151453), the Science & Technology support Program of Suzhou (SYN201503, SNG2017048), and a project funded by the Priority Academic Program Development of Jiangsu Higher Education Institutions. The funders had no role in study design, data collection and analysis, decision to publish, or preparation of the manuscript.

